# Distinct features of multivesicular body-lysosome fusion revealed by a new cell-free content-mixing assay

**DOI:** 10.1101/133074

**Authors:** Mahmoud Abdul Karim, Dieter Ronny Samyn, Sevan Mattie, Christopher Leonard Brett

**Affiliations:** Department of Biology, Concordia University, 7141 Sherbrooke St. W., SP-501.15, Montréal, QC H4R 1R6, Canada; Present address: Montreal Neurological Hospital and Institute, McGill University, Montréal, QC H3A 2B4, Canada; Lead Contact

**Keywords:** SNARE, syntaxin, Pep12, Ypt7, Rab7, Rab-GTPase, multivesicular body, MVB, lysosome, vacuole, membrane fusion, ESCRT, endocytosis, Rab conversion

## Abstract

When marked for degradation, surface receptor and transporter proteins are internalized and delivered to endosomes where they are packaged into intralumenal vesicles (ILVs). Many rounds of ILV formation create multivesicular bodies (MVBs) that fuse with lysosomes exposing ILVs to hydrolases for catabolism. Despite being critical for protein degradation, the molecular underpinnings of MVB-lysosome fusion remain unclear, although machinery underlying other lysosome fusion events is implicated. But how then is specificity conferred? And how is MVB maturation and fusion coordinated for efficient protein degradation? To address these questions, we developed a cell-free MVB-lysosome fusion assay using *S. cerevisiae* as a model. After confirming that the Rab7 ortholog Ypt7 and the multisubunit tethering complex HOPS are required, we found that the Qa-SNARE Pep12 distinguishes this event from homotypic lysosome fusion. Mutations that impair MVB maturation block fusion by preventing Ypt7 activation, confirming that a Rab-cascade mechanism harmonizes MVB maturation with lysosome fusion.

**IMPACT STATEMENT:** Endocytosis culminates with multivesicular bodies (MVBs) fusing with lysosomes. But the molecular underpinnings of this event remain unclear. Here, using *S. cerevisiae* as a model, Karim et al. employ a new in vitro assay to show that MVB-lysosome fusion is driven by ESCRT-dependent Rab-GTPase activation and the syntaxin ortholog Pep12, distinguishing it from other lysosome membrane fusion events.

## INTRODUCTION

Endocytosis regulates surface expression levels of polytopic proteins such as transporters and receptors for cellular signaling and survival in all eukaryotic organisms. Proteins destined for degradation are labeled with ubiquitin and cleared from the surface by invagination and scission of the plasma membrane. Through membrane fusion events, newly formed endocytic vesicles deliver cargo proteins to endosomal membranes where they encounter ESCRTs (<u>E</u>ndosomal <u>S</u>orting <u>C</u>omplexes <u>R</u>equired for <u>T</u>ransport) that sorts them into intraluminal <u>v</u>esicles (ILVs). Many rounds of ILV formation produce a <u>m</u>ulti<u>v</u>esicular <u>b</u>ody (MVB; Huotari and Helenius, 2011; Schmidt and Teis, 2012) that when mature fuses with the lysosome exposing ILVs to lumenal hydrolases for catabolism (Futter et al., 1996; Piper and Katzmann, 2007; Luzio et al., 2010). Although critical for surface protein degradation, we still do not understand many aspects of this terminal step of the endocytic pathway in molecular detail.

However, based on results primarily acquired using genetic approaches and microscopy, a model has emerged describing the mechanisms underlying MVB-lysosome fusion that are thought to be conserved in all eukaryotic species (reviewed by Luzio et al., 2010; Balderhaar et al., 2013). MVBs contain two paralogs of almost every component of the fusion machinery (Kümmel and Ungermann, 2014). One set (the Rab5 module) is required for endosome membrane fusion events responsible for anterograde membrane trafficking to the MVB, for surface protein delivery and organelle biogenesis. The second set (Rab7 module) is proposed to mediate MVB-lysosome fusion. However, most components of the latter set also drive homotypic lysosome fusion (Wickner, 2010; Wickner and Rizo, 2017), provoking the question: What distinguishes these two fusion events?

Because the underlying machinery is similar, inferences into the molecular basis of MVB-lysosome fusion have been drawn from detailed knowledge of the homotypic lysosome membrane fusion reaction, largely elucidated by studying *Saccharomyces cerevisiae* and its vacuolar lysosome (or vacuole), as models. Advances in this area were primarily driven by the use of reliable, quantitative cell-free membrane fusion assays to study homotypic vacuole fusion (Haas, 1995; Jun and Wickner, 2007). This powerful biochemical approach has revealed four distinct subreactions required for organelle membrane fusion and the critical players for each that presumably mediate MVB-vacuole fusion as well:

*Priming*, the first subreaction, requires Sec18, an NSF (<u>N</u>-ethylmaleimide-<u>s</u>ensitive factor) ortholog and homohexameric ATPase, and Sec17, an α-SNAP (<u>s</u>oluble <u>N</u>SF <u>a</u>ttachment <u>p</u>rotein) ortholog and protein chaperone, to unravel cis-SNARE (<u>SNA</u>P <u>re</u>ceptor) complexes from previous fusion events to free up individual SNARE proteins for future rounds of membrane fusion (Mayer et al., 1996; Ungermann et al., 1998). As the only α-SNAP and NSF orthologs in *S. cerevisiae,* Sec17 and Sec18 are thought to mediate all SNARE-mediated membrane fusion events in the cell, but their roles in MVB-vacuole fusion have not been investigated. Afterwards (or possibility concomitantly), organellar membranes undergo *tethering*, the second subreaction, whereby apposing membranes make first contact. This requires activation of the Rab-GTPase Ypt7 (a Rab7 ortholog) and interaction with its cognate <u>m</u>ultisubunit tethering <u>c</u>omplex (MTC) called HOPS (<u>ho</u>motypic fusion and vacuole <u>p</u>rotein <u>s</u>orting complex; Eitzen et al., 2000; Seals et al., 2000).

Next, fusogenic proteins (e.g. Sec17, Ypt7, HOPS and SNAREs) and lipids are recruited to the initial contact site where they assemble into an expanding ring at the vertex between adjacent organelles (Wang et al., 2002; Wang et al., 2003; Fratti et al., 2004). Called *docking*, this subreaction also includes the formation of trans-SNARE protein complexes mediated by HOPS (Collins and Wickner, 2007; Starai et al., 2008). Like homotypic lysosome fusion, Ypt7 and HOPS accumulate at contact sites between MVB and vacuole membranes in *S. cerevisiae* (Cabrera et al., 2009). Furthermore, depleting components of HOPS impairs delivery of endocytic cargoes to vacuoles in yeast or lysosomes in human cells (Peterson and Emr, 2001; Wartosh et al., 2015). Thus both Ypt7 and HOPS are implicated in MVB-vacuole fusion where they presumably contribute to tethering and docking (Bugnicourt et al., 2004; Balderhaar et al., 2013; Pols et al., 2013; Numrich and Ungermann, 2014). During the last *fusion* subreaction, trans-SNARE complexes containing 3 Q-SNAREs (Qa, Qb and Qc donated from one membrane) and one R-SNARE (from the apposing membrane) fully zipper driving lipid bilayer merger (Schwartz and Merz, 2009). Two of four SNAREs within the complex that mediates homotypic lysosome fusion are also found on MVB membranes: the Qb-SNARE Vti1 and soluble Qc-SNARE Vam7 (Fischer von Mollard et al., 1997; Gossing et al., 2013). Otherwise, the MVB expresses its own paralogs of the Qa-SNARE, Pep12 in place of Vam3 on the vacuolar lysosome, and the R-SNARE Snc2 in place of Nyv1 (Gerrard et al., 2000; Gurunathan et al., 2000). The SNAREs donated by each organelle to the complex that drives MVB-lysosome fusion are unknown. Thus, it is possible that the MVB donates either Q-SNAREs including Pep12 or the R-SNARE Snc2 to SNARE complexes distinguishing MVB-lysosome from homotypic lysosome fusion.

Recognizing the impact of these cell-free assays have had on our understanding of homotypic lysosome fusion, we developed a similar method to reliably measure MVB-lysosome membrane fusion in *S. cerevisiae.* Here, we use it to test prevailing model describing the mechanisms underlying this heterotypic fusion event and, more importantly, to uncover mechanisms that differ from homotypic lysosome fusion. One intriguing and unique feature of MVB-lysosome fusion is the need for MVB maturation prior to fusion for efficient surface protein degradation. But how does the mature MVB know when to fuse? Deleting components of ESCRTs block protein sorting into ILVs and impair ILV formation, preventing delivery of internalized proteins to lysosomes and causing endomembrane accumulation. Although implied, there is no empirical evidence supporting the hypothesis that knocking out ESCRTs block this fusion event. Furthermore, how ESCRTs may target the underlying machinery is not entirely understood. Thus, here we use our new fusion assay to better understand how MVB maturation is coupled to lysosome fusion to ensure efficient surface protein degradation.

## RESULTS

### A new cell-free assay to measure MVB-vacuole membrane fusion

We designed our new cell-free MVB-vacuole membrane fusion assay based on a strategy originally devised by Jun and Wickner (2007) to study homotypic vacuole fusion that relies on the reconstitution of β-lactamase upon luminal content mixing (Figure 1A): To target the first fusion probe to the MVB lumen, we fused the C-terminus of the endosomal Qa-SNARE Pep12 to the proto-oncogene product c-Fos followed by the ω-subunit of β-lactamase (Pep12-Fos-Gs-ω). To target our second fusion probe to the vacuole lumen, we fused the targeting sequence of lysosomal protease carboxypeptidase Y (CPY; first 50 amino acids) to Jun, a cognate binding partner of c-Fos, followed by the α-subunit of β-lactamase (CPY50-Jun-Gs-α). We then expressed each probe in separate yeast strains (deficient of vacuolar proteases to prevent probe degradation) to prevent probe interaction in vivo. After isolated organelles from each strain are combined and undergo lipid bilayer merger in vitro, lumenal contents mix allowing Jun and c-Fos to interact driving the complementary halves of β-lactamase together to reconstitute enzyme activity. Reconstituted β-lactamase activity is then measured by monitoring nitrocefin hydrolysis (by recording absorbance at 492 nm over time) to quantify MVB-lysosome fusion.

**Figure 1.**
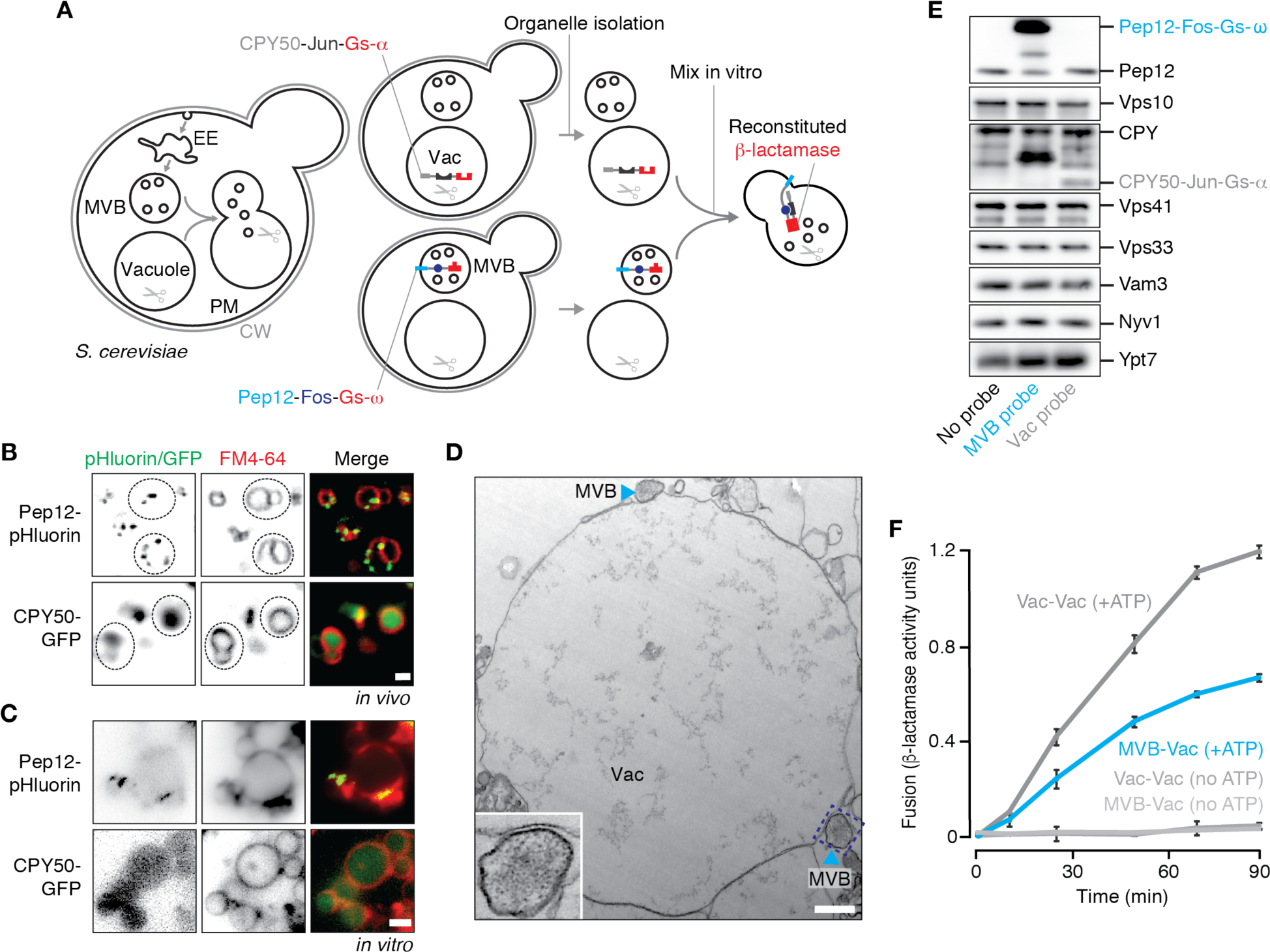
A new cell-free assay to measure MVB-vacuole membrane fusion. (**A**) Cartoon illustrating new cell-free assay to quantify MVB-vacuole fusion. CW, cell wall; PM, plasma membrane; EE, early endosome; Vac, vacuole. Fluorescence micrographs of (**B**) Live cells or (**C**) isolated organelles expressing Pep12-pHluorin or CPY50-GFP. Vacuole membranes are stained with FM4-64. Dotted lines outline yeast cells as observed by DIC. Scale bars, 2 μm. (**D**) Transmission electron micrograph of organelles isolated by ficoll gradient. Structures resembling MVBs are indicated. Insert shows a higher magnification image of the area surrounded by the dotted line. Vac, vacuole. Scale bar, 500 nm. (**E**) Western blots confirming presence of fusion probes and fusogenic proteins in organelles isolated by ficoll gradient. (**F**) Content mixing values obtained over time by mixing organelles isolated from separate strains expressing complimentary fusion probes targeted to MVBs or vacuoles (MVB-Vac), or probes targeted to vacuoles (Vac-Vac), in the presence or absence of ATP to trigger membrane fusion. Means ± S.E.M shown (n ≥ 3).

To confirm that the fusion probes were properly localized within cells, we generated yeast strains expressing versions of the probes tagged with variants of GFP at their C-termini, stained them with FM4-64 to label vacuole membranes and imaged them using fluorescence microscopy (Figure 1B). The Pep12-GFP fusion protein appeared on puncta consistent with MVB localization as reported previously (Becherer et al., 1996), whereas the CPY50-GFP fusion protein was found within the vacuole lumen. Next, we fractionated cellular membranes by sucrose gradient (Figure S1A) and used western blot analysis to confirm that Pep12-Fos-Gs-ω is found in similar fractions (9-12) as Vps10, a resident protein on MVBs (Marcusson et al., 1994), as well as endogenous Pep12. CPY50-Jun-Gs-α is not found in fractions containing the MVB fusion probe, but rather in fractions (3-4) containing Nyv1, a R-SNARE that is exclusively found on vacuole membranes (Wen et al., 2006), and endogenous CPY, a vacuole resident protein. We next collected fractions containing MVB membranes (9-12) from Pep12-Fos-Gs-ω expressing cells or vacuole membranes (3-4) from CPY50-Jun-Gs-α expressing cells and mixed them together under conditions that promote the homotypic vacuole membrane fusion reaction in vitro (see Jun and Wickner, 2007). Unfortunately these preparations were fusion incompetent (Figure S1B). Thus, we sought an alternative method to isolate the organelles to optimize our cell-free fusion assay.

Based on a method to isolate intact, fusogenic vacuoles from yeast cells (see Conradt et al., 1992), we decided to use a 4-step ficoll gradient to isolate both MVBs and vacuoles in the same preparations. Although vacuoles are known to accumulate at the 0-4 % ficoll interface, it is not clear whether MVBs also migrate to this layer during centrifugation. Thus, we performed ficoll-based fractionation experiments using cells expressing GFP-tagged versions of the fusion probes, collected organelles that accumulated at the 0-4% ficoll interface, and imaged them by fluorescence microscopy (Figure 1C). Pep12-GFP is present in the preparation and localizes to small puncta adjacent to vacuole membranes reminiscent of MVB structures, whereas CPY50-GFP is found within the lumen of isolated vacuoles, consistent with their distributions in living cells. We then imaged these organelle preparations using transmission electron microscopy and observed structures resembling MVBs, some of which were in close contact with vacuole membranes, a prerequisite for fusion (Figure 1D; see Mattie et al., 2017). We also observed the presence of our fusion probes (Pep12-Fos-Gs-ω or CPY50-Jun-Gs-α) in this fraction by western blot analysis (Figure 1E). Markers of both MVBs (Vps10, endogenous Pep12) and vacuoles (Nyv1, endogenous CPY), and fusogenic proteins (e.g. Ypt7, Vps41, Vps33 and Nyv1) were also present confirming that both organelles are found in this preparation and are likely fusogenic.

We next mixed equal proportions of organelles isolated from cells expressing only the MVB fusion probe or only the vacuole fusion probe in fusion buffer that mimics cytosolic ^conditions (125 mM KCl, 5 mM MgCl2, 200 mM sorbitol, 20 mM PIPES, pH 6.80). We added 1^ mM ATP to trigger membrane fusion and incubated reactions at 27˚C for up to 90 minutes. Upon measuring reconstituted β-lactamase activity (Figure 1F), we observed robust nitrocefin hydrolysis indicative of heterotypic fusion, showing a signal up to 10-fold greater than background (i.e. reactions run without ATP or left on ice). Rates of heterotypic fusion were comparable to those observed for homotypic vacuole fusion, an additional indication that this new assay may be reliably used to study the heterotypic fusion reaction in detail.

### Sec17, Ypt7 and HOPS drive MVB-vacuole fusion in vitro

With this new assay in hand, we next tested the prevailing model of MVB-vacuole fusion (Figure 2A), which predicts that Sec17, Ypt7 and HOPS orchestrate this event (Balderhaar et al., 2010, 2013). The α-SNAP ortholog Sec17 both unravels cis-SNARE complexes and, through HOPS binding, promotes SNARE-zippering during homotypic fusion (Schwartz and Merz, 2009; Wickner and Rizo, 2017). As the only α-SNAP ortholog in *S. cerevisiae*, it is hypothesized to also play a role in MVB-vacuole fusion. Indeed, adding purified anti-Sec17 antibody, at concentrations that block homotypic fusion, also prevents heterotypic fusion (Figure 2B), suggesting a role in both events. We next confirmed that Ypt7 and HOPS (the Rab7 module in yeast) also underlie both fusion events (Figure 2B): Addition of the Rab-GTPase inhibitors rGdi1 (a Rab-GTPase chaperone that extracts Rab proteins from membranes) or rGyp1-46 (the active domain of the Rab-GAP protein Gyp1; see Brett and Merz, 2008) blocks both fusion events, confirming that Rab-GTPase activation is required. Addition of purified anti-Ypt7 antibody, at levels that block homotypic fusion, inhibited heterotypic fusion, confirming that activation of this specific Rab is needed for both fusion events. The HOPS holocomplex contains six protein subunits, two of which are unique (Vps41 and Vps39) and four (Vps33, Vps11, Vps18 and Vps16) that are shared with CORVET (class <u>C</u> c<u>or</u>e <u>v</u>acuole/<u>e</u>ndosome tethering complex), a similar multisubunit tethering complex within the Rab5 module also found on endosomes (Peplowska et al., 2007; Nickerson et al., 2009; Plemel et al., 2012). Addition of purified anti-Vps33 antibody blocks both fusion events (Figure 2B), suggesting involvement of either complex. Purified anti-Vps41 antibody also blocked heterotypic and homotypic fusion (Figure 2B), confirming that HOPS, specifically, contributes MVB-vacuole fusion.

**Figure 2.**
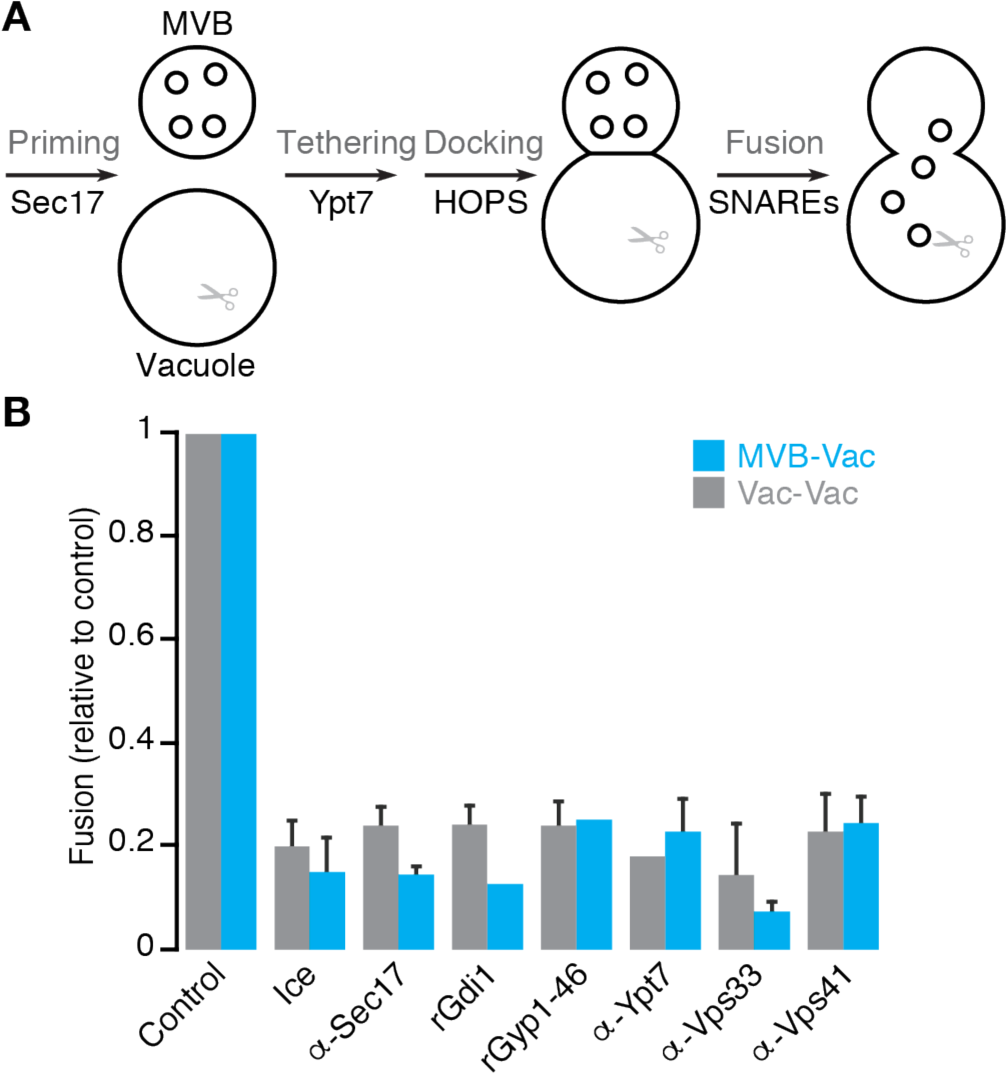
Priming, tethering and docking machinery is shared by hetero- and homo-typic fusion events. (**A**) Cartoon illustrating the predominant model of MVB-vacuole fusion. Proteins or protein complexes implicated in subreactions are indicated. (**B**) MVB-vacuole (MVB-Vac) or homotypic vacuole (Vac-Vac) fusion measured in the presence or absence of affinity-purified antibodies against Sec17 (1.8 μM), Ypt7 (1.8 μM), Vps33 (1.8 μM) or Vps41 (1.8 μM), or purified recombinant Gdi1 (4 μM) or Gyp1-46 (5 μM) proteins. Reactions were kept on ice to prevent fusion as negative controls. Means ± S.E.M. shown (n ≥ 3).

### Trans-SNARE complexes distinguish MVB-vacuole from homotypic vacuole fusion events

The final stage of the organelle fusion reaction is lipid bilayer merger driven by complete trans-SNARE-complex zippering (Nichols et al., 1997). For homotypic vacuole fusion, trans-SNARE complexes are minimally composed of the three Q-SNAREs – the syntaxin ortholog Vam3 (Qa), Vti1 (Qb) and the SNAP25 ortholog Vam7 (Qc) – and the R-SNARE Nyv1, a synaptobrevin ortholog (Wickner, 2010; Wickner and Rizo, 2017). Vam7, a soluble protein, complexes with Vam3 and Vti1 on one membrane forming a Q-SNARE bundle that binds to Nyv1 on the apposing membrane (Schwartz and Merz, 2009). Of these 4 SNAREs, only Vti1 and Vam7 are expressed on MVBs. Otherwise MVBs express the Qa-SNARE paralog Pep12 and R-SNARE Snc2 (Becherer et al., 1996; Fischer von Mollard et al., 1997; Gerrard et al., 2000; Gurunathan et al., 2000) suggesting that trans-SNARE complexes mediating heterotypic and homotypic fusion are different, whereby the MVB donates either Pep12 or Snc2 (but not both). Elegant work by Joji Mima’s group shows that synthetic proteoliposomes containing Pep12, in place of the vacuolar Qa-SNARE Vam3, are capable of forming stable, fusogenic complexes with the other SNAREs responsible for homotypic vacuole fusion (Furukawa and Mima, 2014). Thus, we hypothesized that a trans-SNARE complex composed of Pep12, Vti1 and Vam7 donated from MVBs with Nyv1 donated by vacuoles may drive heterotypic fusion events.

To test this hypothesis, we isolated SNARE complexes using an approach that ensures they form in trans during the membrane fusion reaction in vitro (see Collins and Wickner, 2007; Schwartz and Merz, 2009): We mixed organelles missing NVY1 and containing Pep12 fused to calmodulin-binding protein (CBP::Pep12) with organelles missing PEP12 under fusogenic conditions, and then conducted pull-down assays and western blot analysis to identify SNARE proteins in complex with CBP::Pep12 (Figure 3A). We also repeated the experiment with CBP::Vam3 for comparison and as a positive control (Schwartz and Merz, 2009). As expected, both trans-SNARE complexes contained Vti1 and Vam7, but only under fusogenic conditions (with ATP; Figure 3B). Very little Vma3 co-purified with CBP::Pep12, and no Pep12 was found in CBP::Vam3 complexes, confirming the presence of distinct Q-SNARE bundles. Importantly, the vacuolar R-SNARE Nyv1 was found in both complexes, but not the endosomal R-SNARE Snc2, confirming that Pep12-Vti1-Vam7-Nyv1 complexes form in trans during organelle fusion in vitro.

**Figure 3.**
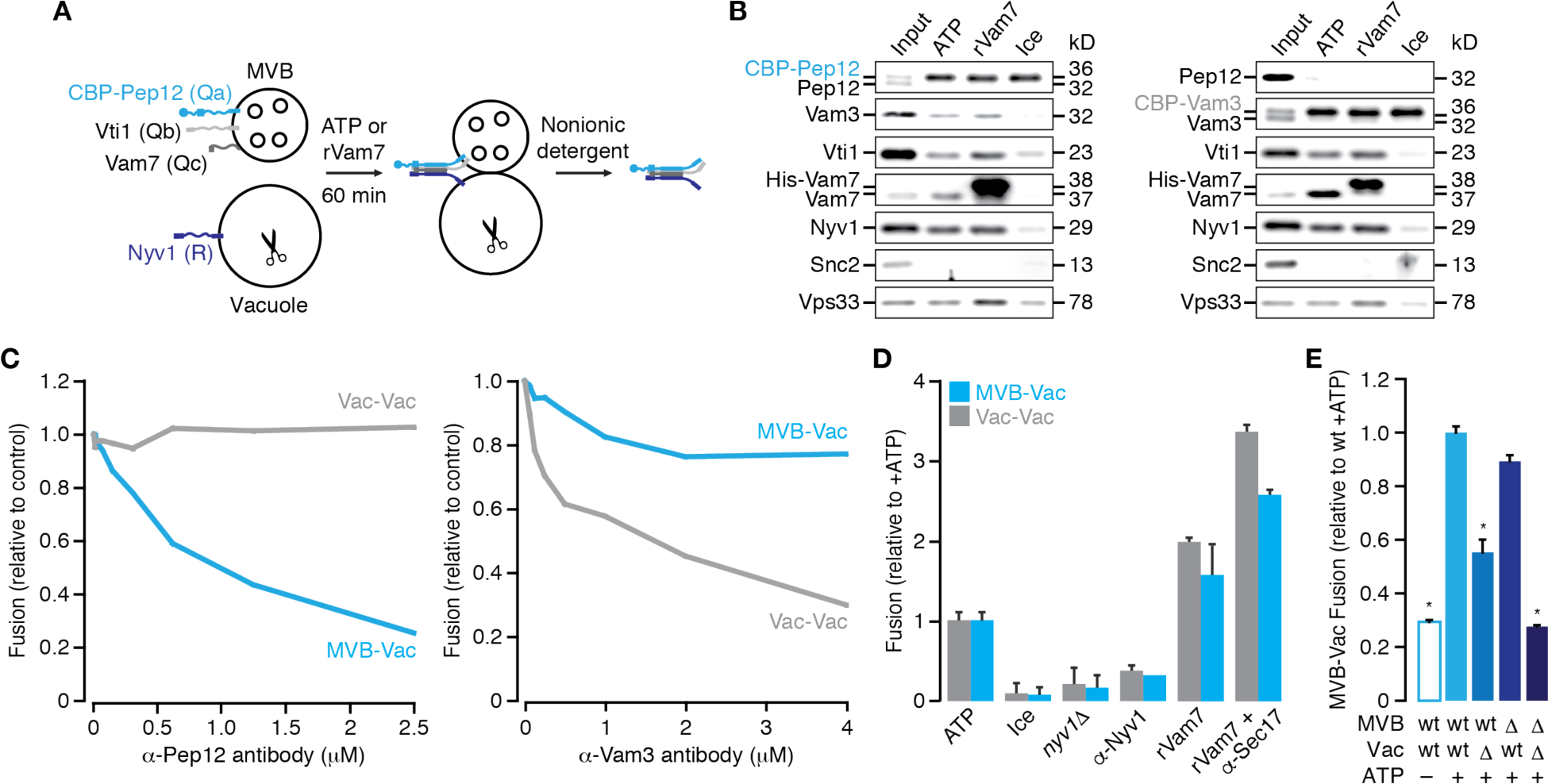
Trans-SNARE complexes distinguish vacuole fusion events. (**A**) Cartoon illustrating method used to isolate trans-SNARE complexes from organelle fusion reactions in vitro. CBP-Vam3 was used in place of CBP-Pep12 to isolate SNARE complexes known to mediate homotypic vacuole fusion. (**B**) Western blots used to identify fusogenic proteins bound to CBP-Pep12 (left) or CBP-Vam3 (right) based on pull-down experiments illustrated in **A**. (**C**) MVB-vacuole (MVB-Vac) or homotypic vacuole (Vac-Vac) fusion measured in the presence of increasing concentrations of purified anti-Pep12 (left) or anti-Vam3 (right) antibody. (**D**) ATP-driven heterotypic (MVB-Vac) or homotypic (Vac-Vac) fusion measured using organelles isolated from wild type or NYV1 knock out cells (*nyv1*∆) or from wild type cells in the absence or presence of 2.2 µM purified anti-Nyv1 antibody. Fusion of organelles from wild type cells was also measured in the presence of 100 nM rVam7 to stimulate fusion in place of ATP, with or without pretreatment with 1.8 µM anti-Sec17 to encourage SNARE complex formation. Reactions were kept on ice to prevent fusion as negative controls. (**E**) Organelles isolated from wild type (wt) or NYV1 knock out (∆) cells were mixed in vitro and MVB-vacuole fusion was measured in the presence or absence of ATP. *P < 0.05, as compared to value obtained with only organelles from wild type cells in the presence of ATP. Means ± S.E.M. shown (n ≥ 3).

Addition of purified, recombinant Vam7 protein (rVam7, the soluble Qc-SNARE) to isolated vacuoles drives formation vacuolar trans-SNARE complexes promoting homotypic fusion without ATP (Thorngren et al., 2004). Given that Vam7 is found in both trans-SNARE complexes, we next tested if rVam7 could also stimulate assembly of Pep12-containing complexes. As expected, rVam7 drives formation of both trans-SNARE complexes (Figure 3B), suggesting that it contributes to heterotypic as well as homotypic fusion. Vps33, a component of HOPS and SM-protein ortholog, binds SNAREs and initiates trans-SNARE complex formation during homotypic vacuole fusion by threading the Qa-SNARE Vam3 with the R-SNARE Nyv1 (Baker et al., 2015). Recombinant Vps33 protein was also shown to thread Pep12 and Nyv1 proteins in vitro (Lobingier et al., 2012). Thus, we also tested whether Vps33 binds to trans-SNARE complexes thought to mediate MVB-vacuole fusion. Indeed, Vps33 bound to Pep12-Nyv1-containing as well as Vam3-Nyv1-contianing complexes (Figure 3B), suggesting that HOPS initiates formation of Pep12-Vti1-Vam7-Nyv1 complexes in trans to drive MVB-vacuole fusion.

To demonstrate that this distinct trans-SNARE complex drives MVB-vacuole fusion, we next tested the role of these SNAREs in this process using our new in vitro assay. First, we added different purified antibodies raised against the Qa-SNAREs Pep12 or Vam3 to organelle fusion reactions (Figure 3C). As predicted, increasing concentrations of anti-Pep12 antibody preferentially blocked heterotypic fusion, whereas anti-Vam3 antibody preferentially inhibited homotypic fusion. Addition of purified anti-Nyv1 antibody or knocking out NYV1 blocked both fusion events, consistent with the presence of this R-SNARE in both complexes (Figures 3D and S2). Moreover, when we mix mutant and wild type organelles, we find that heterotypic fusion is impaired only when vacuole membranes lack NVY1 (Figure 3E), confirming that this R-SNARE is donated from the vacuole. To confirm that trans-SNARE complexes formed by the Qc-SNARE Vam7 mediate both MVB-vacuole and homotypic vacuole fusion, we next added rVam7 to isolated organelles in the absence of ATP (Figure 3D). As for homotypic fusion (Thorngren et al., 2004), we found that addition of rVam7 is sufficient to stimulate MVB-vacuole fusion in vitro. Furthermore, addition of anti-Sec17 antibody to prevent unraveling of newly formed SNARE complexes (Schwatrz and Merz, 2009) enhanced this effect, confirming that it contributes to heterotypic as well as homotypic fusion. Together, these results demonstrate that the Pep12-Vti1-Vam7-Nyv1 trans-SNARE complex drives MVB-vacuole fusion, revealing that the composition of SNARE complexes distinguishes heterotypic from homotypic fusion events.

### ESCRTs activate Ypt7 for MVB-lysosome fusion

The ESCRT machinery mediates sorting and packaging of internalized surface proteins into intralumenal vesicles creating the MVB (Henne et al., 2011). Deleting components of ESCRTs prevents delivery of internalized surface proteins to vacuoles and causes them to accumulate on aberrant, enlarged endocytic compartments (Raymond et al., 1992; Robinson et al., 1998; Russel et al., 2012). These observations led to the hypothesis that protein sorting and ILV formation by ESCRTs is necessary to trigger MVB-lysosome fusion (Metcalf and Issacs, 2010). To directly test this hypothesis, we individually knocked out single components of three ESCRTs (VPS23 in ESCRT-I, VPS36 in ESCRT-II and SNF7 in ESCRT-III; Henne et al., 2011) in strains expressing the MVB fusion probe. After confirming the probe localized to aberrant endocytic compartments within these mutant cells (Figure 4A; see Coonrod and Stevens, 2010; Russell et al., 2012), we mixed organelles isolated from these strains with organelles from wild type cells expressing the vacuole fusion probe and conducted in vitro fusion assays (Figure 4B). This approach ensured that vacuoles were fusogenic, as deleting components of ESCRTs blocks the CPY and CPS (carboxypeptidase-S) biosynthetic pathways that supply vacuoles with fusion proteins (Katzmann et al., 2001) – noting that this also prevented us from assessing effects of these mutants on homotypic vacuole fusion in vitro. As predicted, deleting any of the three ESCRT components inhibited MVB-vacuole fusion, indicating that blocking any stage of protein sorting into ILVs impairs heterotypic fusion.

**Figure 4.**
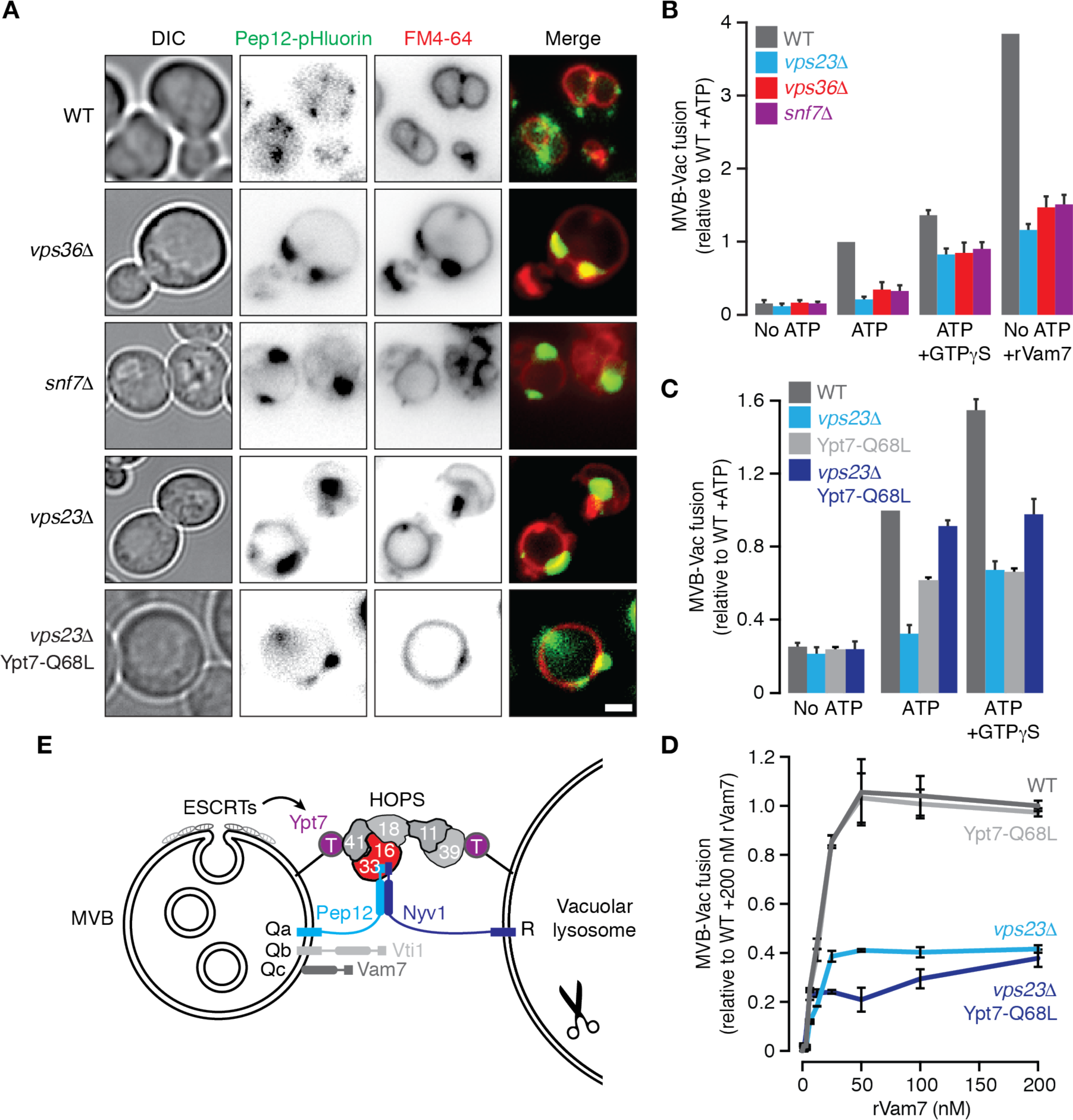
Impaired ESCRT function prevents Ypt7 activation and MVB-vacuole fusion. (**A**) Fluorescence micrographs of live wild type (WT), *vps36*∆, *snf7*∆, *vps23*∆ or *vps23*∆ Ypt7-Q68L cells expressing Pep12-pHluorin. Vacuoles membranes were stained with FM4-64. Scale bar, 2 μm. (**B**) Fusion values from reactions containing organelles isolated from WT, *vps23*∆, *vps36*∆ or *snf7*∆ cells expressing the MVB fusion probe mixed with organelles isolated from wild type cells expressing the vacuole fusion probe in the absence or presence of ATP, 0.2 mM GTPγS, or 100 nM rVam7. (**C,D**) Fusion values from reactions containing organelles isolated from WT, *vps23*∆, Ypt7-Q68L or *vps23*∆ Ypt7-Q68L cells expressing the MVB fusion probe mixed with organelles isolated from wild type cells expressing the vacuole fusion probe in the absence or presence of ATP or 0.2 mM GTPγS. (**C**) or increasing concentrations of recombinant Vam7 protein (**D**). (**E**) Cartoon illustrating new model of MVB-vacuole fusion. Means ± S.E.M. shown (n ≥ 3).

How do ESCRTs stimulate MVB-vacuole fusion? Rab-GTPase conversion coordinates successive fusion events in the secretory and endocytic pathways for efficient delivery of cargo proteins (Rivera-Molina and Novick, 2009). At the MVB, it has been proposed that inactivation of the Rab5 ortholog Vps21 (responsible for endosomal fusion events that deliver proteins to the MVB) is necessary for activation of Ypt7 to initiate the next trafficking step: MVB-vacuole fusion. Impairing protein sorting into ILVs by ESCRTs causes active Vps21 to accumulate on aberrant enlarged endosomes (Russell et al., 2012). Thus, it was hypothesized that deleting ESCRTs impairs Rab-conversion, whereby Ypt7 cannot be activated to trigger MVB-vacuole fusion. To test this hypothesis, we took three approaches:

First, we added GTPγS, a non-hydrolyzable GTP analog that activates all GTPases (including Ypt7) at a concentration (0.2 mM) that supports vacuole-vacuole and MVB-vacuole fusion in vitro (Jun and Wickner, 2007; Mattie et al., 2017; Karim and Brett, 2017), and found that it partially rescued heterotypic fusion defects caused by deleting components of ESCRTs (Figure 4B). Next, we added 100 nM rVam7 in place of ATP to stimulate fusion, as it bypasses the requirement for Ypt7 (Thorngren et al., 2004), and found that it also partially rescued fusion defects (Figure 4B and D). These results also confirm that Ypt7 and SNAREs are functional, but not engaged, on aberrant endosomal compartments isolated from ESCRT mutants. Finally, to demonstrate that Ypt7 is targeted, we introduced a Q68L mutation in YPT7 to render it constitutively active (see Brett et al., 2008) in cells expressing the MVB fusion probe and lack VPS23. After confirming the fusion probe was properly localized, we found that it too rescued MVB-lysosome fusion defects caused by this mutation (Figures 4A and 4C). Addition of GTPγS did not further enhance heterotypic fusion in the presence of Ypt7 Q68L, suggesting that Ypt7 on the MVB membrane was the likely target of GTPγS that rescues fusion when ESCRT function is impaired. Furthermore, the mutation did not affect rVam7-driven fusion in the presence or absence of VPS23 (Figure 4D), confirming that Rab activity is bypassed under these conditions. In all, these findings demonstrate that disrupting the ESCRT machinery prevents Ytp7 activation on MVB membranes required for MVB-vacuole fusion.

## DISCUSSION

### A reliable cell-free assay to study MVB-lysosome fusion

Here we describe a new cell-free assay to measure MVB-vacuole membrane fusion based on reconstitution of lumenal β-lactamase (Figure 1). The method of organelle isolation was crucial for success, and although MVBs are not separated from vacuoles in these preparations, targeting the fusion probes to each organelle in separate yeast strains and mixing them later ensures that reconstituted β-lactamase activity represents only fusion that occurs in vitro under controlled conditions. Importantly, this new assay offers a reliable method to study the MVB-vacuole membrane fusion reaction in detail, by providing many experimental advantages:

First, it is quantitative, robust (with a signal-to-noise ratio up to 10:1) and allows kinetic analysis. Second, it is a simple, colorimetric alternative to fluorescence-based assays limited by resolution and accuracy of colocalization methods offered by light microscopy (offering only 3:1 signal-to-noise; Cao et al., 2015) or assays that require use of radioactive reagents, e.g. ^125^I-labeled biotin (offering only 4:1 signal-to-noise; Pryor et al., 2004). Third, it overcomes limitations associated with using genetic approaches to inhibit potential contributors to this event, whereby mutations hypothesized to block MVB-vacuole fusion also impair carboxypeptidase-Y and -S biosynthetic pathways that feed resident proteins to vacuoles, possibly introducing defects that affect fusogenicity in vivo (see Robinson et al., 1988; Raymond et al., 1992). This is achieved by adding protein or small molecule inhibitors directly to fusion reactions containing organelles isolated from wild type cells or by mixing organelles isolated from mutants and wild type cells expressing complimentary probes; an approach that may also be used to define contributing mechanisms from each organelle. (4) It overcomes the need to purify organelles to homogeneity, as required to make inferences from alternative methods that do not selectively label a single organelle population, e.g. lipid dyes (R-18) for fluorescence dequenching-based lipid-mixing fusion assays (e.g. Jun and Wickner, 2007), or internalized fluorescence dyes that label multiple endocytic compartments (e.g. Morvan et al., 2009). Fifth, unlike most cell-free endosome or MVB fusion assays (e.g. Vida and Gerhardt, 1999), it does not require addition of cytosol indicating that all machinery necessary for fusion co-purifies with organelles facilitating their study. Finally, it offers the advantages of changing the reaction buffer conditions to simulate changes in the “cytoplasm” to better understand how heterotypic fusion is regulated or using protein reagents to overcome issues associated with pleiotropy, allowing study of contributions from gene products predicted to have multiple functions at multiple sites in cells, e.g. Sec17.

### A new model describing MVB-lysosome fusion

Here we definitively show that Sec17, Ypt7 and the HOPS complex are necessary for membrane fusion between MVBs and vacuoles (Figure 2). These proteins also contribute to other fusion events in cells (Balderhaar et al., 2013; Kümmel and Ungermann, 2014). Thus, deleting the genes encoding them induces pleiotropic phenotypes, including blockade of biosynthetic pathways to the vacuole (Robinson et al., 1988; Peterson and Emr, 2001). This work therefore lends important biochemical-based evidence to support the prevailing model describing the molecular underpinnings of MVB-vacuole fusion (Figures 2A and 4E), which was almost exclusively backed by evidence generated using genetic approaches (Kümmel and Ungermann, 2014). Moreover, we make an important advance in our understanding of this process:

A unique trans-SNARE complex distinguishes MVB-vacuole fusion from other vacuole fusion events. It differs from the complex that drives homotypic vacuole fusion whereby the Qa-SNARE Pep12 is donated from the MVB, in place of the vacuolar paralog Vam3, to form a Pep12-Vti1-Vam7-Nyv1 complex in trans (Figure 3B). As preassembled Q-SNARE bundles are donated from one membrane and R-SNAREs from the other (Schwartz and Merz, 2009), we propose that the MVB donates the Q-SNARE bundle Pep12-Vti1-Vam7 and the vacuole donates the R-SNARE Nyv1. We also demonstrate that these SNAREs are necessary or sufficient for MVB-vacuole fusion using three approaches (Figure 3C–E): deleting NYV1 from vacuole membranes blocks fusion, adding purified antibodies raised against Nyv1 or Pep12 blocks fusion, or adding purified, recombinant Vam7 protein drives fusion. Importantly, we measure homotypic vacuole fusion as a control (Figures 1 – 3) to demonstrate that reagents used to target shared machinery are effective at the concentrations shown. We also exclude the possibility that the inverse trans-SNARE complex arrangement contributes to this process, i.e. a complex containing the Qa-SNARE Vam3 (in complex with Vti1 and Vam7) from the vacuole and R-SNARE Snc2 donated by the MVB (Figure 3). These findings are consistent with numerous previous reports:

First, when Pep12-Vti1-Vam7-containing synthetic proteoliposomes are mixed with proteoliposomes expressing Nyv1, these SNAREs form stable protein complexes in trans that support lipid bilayer fusion (Furukawa and Mima, 2014). Second, Vam3 contributes to the ALP (alkaline phosphatase) biosynthetic pathway, which sends newly synthesized proteins directly from the Golgi to the vacuole bypassing the endosome or MVB (Cowles et al., 1997). Pep12, on the other hand, is implicated in the CPY biosynthetic pathway, which sends proteins from the Golgi to endosomes or MVBs en route to the vacuole (Becherer et al., 1996; Gerrard et al., 2000), consistent with this Qa-SNARE, and not Vam3, mediating MVB-vacuole fusion. Third, overexpression of VAM3 suppresses membrane trafficking defects in *pep12*∆ yeast cells (Götte and Gallwitz, 1997), suggesting that Pep12 and Vam3 can be exchanged within SNARE complexes as we show in Figure 3B. Forth, consistent with our findings, Q-SNAREs donated by the endosome membrane and an R-SNARE donated by the lysosome membrane are proposed to mediate endosome-lysosome fusion in mammalian cells (Pryor et al., 2004), suggesting this mechanism is evolutionarily conserved. Finally, Vps33 is known to function at the endosome and vacuole (Subramanian, 2004) and studies conducted with purified proteins demonstrate that the SM-protein Vps33 binds to Vam3 or Pep12 with equal affinity to promote templating with Nyv1 (Lobingier and Merz, 2012; Baker et al., 2015). Here we show that Vps33 binds SNARE complexes containing Pep12-Nyv1 or Vam3-Nyv1 (Figure 3B) and MVB-vacuole fusion requires Vps33 and Pep12 but not Vam3 (Figures 2B and 3C). Thus, we conclude that Vps33 within the HOPS complex promotes Pep12-Vti1-Vam7-Nyv1 complex formation in trans to drive MVB-lysosome fusion (Figure 4E).

However, our findings are not entirely consistent with work published by Stevens and colleagues who show that deleting NYV1 does not affect CPY delivery to the vacuole lumen which presumably requires MVB-vacuole fusion (Fischer von Mollard and Stevens, 1999; also see Figure S2). One possible explanation is that Ykt6, a lipid-anchored R-SNARE also found on vacuole membranes, compensates for the loss of Nyv1 by replacing it in SNARE complexes that drive this fusion event in living cells (Ungermann et al., 1999). However, Ykt6 dissociates from isolated vacuole membranes (*Dietrich et al., 2005*), possibly explaining why it does not replace Nyv1 to drive MVB-vacuole fusion in vitro. Furthermore, it is not clear whether Ykt6 efficiently complexes with Qa-SNARE bundles containing Pep12 or contributes to MVB-vacuole fusion. As far as we know, reliable antibodies raised against Ykt6 are unavailable and thus, we were unable to comprehensively test this hypothesis. However, we plan to address this issue in future studies that will determine if Ykt6 instead plays a role in fusion between vacuoles and other organelles.

### ESCRTs trigger MVB-lysosome fusion by promoting Rab conversion

Although they perform unique roles in polytopic protein sorting and packaging for degradation, deleting any ESCRT component causes formation of aberrant, enlarged endosomal structures called “class E compartments” to appear instead of proper MVBs (e.g. Russell et al., 2012). Here, we show that individually deleting VPS23, VPS326 or SNF7–– components of ESCRT-I, -II and –III, respectively––prevents fusion between these class E compartments and vacuoles in vitro (Figure 4B), consistent with observations that internalized surface proteins or biosynthetic cargoes (e.g. CPY) are not efficiently delivered to vacuoles and accumulate on these aberrant structures (Stuffers et al., 2009). Activating Ypt7 by introducing a Q68L mutation on the class E compartment membrane or adding GTPγS to reactions partially rescued fusion defects caused by deleting these ESCRT components (Figure 4B and C). Although it overcomes defects to support membrane fusion, it is unlikely that Ypt7Q68L allows recovery of protein sorting into ILVs or ILV formation in the absence of VPS23, because it does not play a role in this process. We therefore reason that protein sorting into ILVs and ILV formation act upstream to promote Ypt7 activation on mature MVB membranes, which in turn triggers fusion with vacuoles (Russell et al., 2012). But how does this occur?

The precise molecular mechanism remains unknown, however previous work revealed that active Vps21 and its GEF (<u>G</u>uanine <u>E</u>xchange <u>F</u>actor) Vps9 accumulate on aberrant endosome membranes in cells lacking ESCRT function (Russell et al., 2012). Although Ypt7 is present, its effector Vps41 (and presumably the HOPS complex) is absent from these compartments suggesting Ypt7 is not activated. These findings led to the hypothesis that a Rab conversion mechanism may couple MVB maturation with vacuole fusion, whereby upon completion of ESCRT-dependent protein sorting into ILVs, Vps21 is inactivated, which in turn triggers activation of Ypt7 to promote fusion of the mature MVB with lysosomes. Vps9 is recruited to endosomal membranes by binding ubiquitin linked to internalized surface proteins (Shideler et al., 2015). Normally these proteins are cleared from the perimeter membrane by ESCRTs, causing dissociation of Vps9 from the perimeter membrane and, in turn, Vps21 is inactivated linking MVB maturation to Rab conversion. However, ESCRT impairment causes ubiquitylated proteins to accumulate, Vps9 then remains and chronic activation of Vps21 prevent Rab conversion thus blocking MVB-vacuole fusion (Russell et al., 2012; Shideler et al., 2015). Although the mechanism that couples Vps21 and Ypt7 activities remains elusive, it likely involves stimulation of Msb3, a GAP that inactivates Vps21 (Lachmann et al., 2012), Mon1-Ccz1, the GEF that activates Ypt7 (Nordmann et al., 2010), and possibly BLOC-1 (<u>b</u>iogenesis of lysosome-like <u>o</u>rganelles <u>c</u>omplex-1) to coordinate GAP and GEF function on mature MVB membranes (Rana et al., 2015). We will test this hypothesis in the near future using this new cell-free MVB-vacuole fusion assay to provide a comprehensive understanding of this process and to improve our knowledge of Rab-conversion, which is thought to coordinate progressive membrane trafficking at most sites within eukaryotic cells.

As with the fusion machinery, ESCRTs and potential mediators of Vps21-Ypt7 conversion are evolutionarily conserved, suggesting the same mechanisms likely underlie MVB-lysosome fusion in metazoan cells (Saksena and Emr, 2009; Dennis et al., 2016). Loss-of-function mutations in ESCRT components are linked to many human disorders, including cancers and neurodegenerative diseases (Saksena and Emr, 2009). Similar to phenotypes observed in *S. cerevisiae*, enlarged, abnormal endosome-like structures accumulate and surface protein sorting and degradation is impaired within human cells harboring ESCRT mutations (Stuffers et al., 2009), suggesting MVB-lysosome fusion is defective. Here we show that stimulating Ypt7, the yeast ortholog of human Rab7, overcomes this fusion defect. Although these compartments lack ILVs, this fusion event delivers internalized surface proteins to the vacuolar lysosome membrane where they could be selectively degraded by the IntraLumenal Fragment (ILF) pathway (McNally and Brett, 2017). As it is proposed that inefficient surface receptor degradation underlies the etiology of these diseases, we speculate that developing therapeutics designed to activate Rab7 may represent a valid strategy for treatment.

## MATERIALS AND METHODS

### Yeast strains and reagents

To prevent degradation, all fusion probes were expressed in BJ35050, a *Saccharomyces cerevisiae* strain devoid of vacuolar lumenal proteases [*MATα pep4::HIS3 prbΔ1-1.6R his3– Δ200 lys2–801trp1-Δ101 (gal3) ura3–52 gal2 can1*]. To ensure high levels of protein expression, BJ3505 cells were transformed with two integrating plasmids each containing a copy of a single fusion probe behind a strong (ADH1) promoter: pRS406-Pep12-Fos-Gs-ω (pCB002) and pRS404-Pep12-Fos-Gs-ω (pCB003) contain MVB-targeted probes, and pRS406-CPY50-Jun-Gs-α (pYJ406-Jun; Jun and Wickner, 2007) and pRS404-CPY50-Jun-Gs-α (pCB011) contain vacuole-targeted probes. For homotypic vacuolar lysosome fusion, plasmids containing MVB probes were replaced with pRS406-CPY50-Fos-Gs-ω (pYJ406-Fos; Jun and Wickner, 2007) and pRS404-CPY50-Fos-Gs-ω (pCB008) to target the Gs-ω fragment of β-lactamase to the vacuole lumen instead. To assess effects of Ypt7-Q68L on fusion, we transformed BJ3505 *YPT7*_*Q68L::neo*_ ells (see Brett et al., 2008) with plasmids encoding MVB-targeted fusion probes. NYV1, YPT7, VPS23, VPS36, and SNF7 were knocked out of fusion strains using the Longtine method (Longtine et al., 1998). To assess the cellular location of fusion probes by fluorescence microscopy, we replaced genes encoding c-Fos/Jun and β-lactamase fragments in the fusion probe plasmids with either GFP in pRS406-CPY50-GFP (pCB044) encoding a vacuole-targeted probe, or with pHluorin (a variant of GFP; see Karim and Brett, 2017) in pRS406-Pep12-pHluorin (pCB046) encoding a MVB-targeted probe. To evaluate trans-SNARE complexes, we used BJ3505 cells lacking NYV1 transformed with pRS406-CBP-Pep12 (pCB045) or BJ3505-CBP-Vam3 *nyv1Δ* cells (a gift from Alexey Merz; see Collins and Wickner, 2007).

Biochemical and yeast growth reagents were purchased from Sigma-Aldrich, Invitrogen Bio-Rad Laboratories (Canada) Ltd or BioShop Canada Inc. Restriction enzymes, DNA polymerases and ligases were purchased from New England Biolabs (Ipswich, MA, USA). Calmodulin Sepharose 4B, Ni-Sepharose 6FF and Glutathione Sepharose 4B beads were purchased from GE Healthcare (Missassauga, ON, Canada). Purified rabbit polyclonal antibodies raised against components of the fusion machinery were generous gifts from William Wickner (Dartmouth College, Hanover, NH, USA; Sec17), Alexey Merz (University of Washington, Seattle, WA, USA; Vam3, Vam7, Ypt7, Vps21, Vps33, Vps41, Vps10 and CPY) and Jeffrey Gerst (Weizmann Institute of Science, Rehovot, Israel; Snc2). Recombinant mouse antibody raised against Pep12 was purchased from Abcam (Toronto, ON, Canada). Recombinant Gdi1 protein was purified from bacterial cells using a calmodulin-binding peptide intein fusion system (see Brett and Merz, 2008). Recombinant Gyp1-46 protein (the catalytic domain of the Rab-GTPase activating protein Gyp1) and recombinant Vam7 were purified as previously described (Eitzen et al., 2000; Schwartz and Merz, 2009; respectively). Reagents used in fusion reactions were prepared in 10 mM Pipes-KOH, pH 6.8, and 200 mM sorbitol (Pipes-sorbitol buffer, PS).

### Membrane fractionation by sucrose gradient

Yeast cells were grown in 1L nutrient-rich growth media (YPD) overnight to OD600_nm_ = 1.6/ml, harvested, and treated with lyticase for 30 minutes at 30°C. Spheroplasts were then sedimented and resuspended in 10 ml of TEA buffer (10 mM triethanolamine, pH 7.5, 100 mg/ml phenylmethylsulfonyl fluoride, 10 mM NaF, 10 mM NaN_3_, 1 mM EDTA, and 0.8 M sorbitol) and Dounce homogenized on ice (20 strokes). Lysates were then centrifuged at 15,000 g for 20 minutes, and the resulting supernatant was centrifuged at 100,000 g for 2 hours to sediment cell membranes. Pellets were resuspended in 1 ml TEA buffer and loaded onto a stepwise (20–70%) sucrose density gradient, and then centrifuged at 100,000 g for 16 hours at 4°C to separate membranes by density. Samples were collected from the top, and each fraction was precipitated using 10% trichloroacetic acid, washed and resuspended in 100 μl SDS-PAGE buffer. Fractions were then loaded into SDS-polyacrylamide gels and subjected to electrophoresis to separate proteins by size. Western blot analysis was performed to determine fractions that contained proteins of interest.

### Organelle isolation and cell-free fusion assay

Organelles were isolated from yeast cells by ficoll floatation, as previously described (Haas, 1995). Organelle membrane fusion was assessed using a modified version of a lumenal content mixing assay that relies on reconstitution of β-lactamase activity (Jun and Wickner, 2007). In brief, organelles were isolated from two separate strains that each express a single complementary fusion probe, and 6 μg of each were added to 60 μl reactions containing standard fusion buffer (125 mM KCl, 5 mM MgCl_2_, 1 mM ATP, 40 mM creatine phosphate, 0.5 mg/ml creatine kinase, and 10 μM CoA in PS buffer) supplemented with 10 μM recombinant GST-Fos protein to reduce background caused by organelle lysis. Reactions were incubated up to 90 minutes at 27°C and then stopped by placing them on ice. Lumenal content mixing was then quantified by measuring the rate of nitrocefin hydrolysis by reconstituted β-lactamase: 58 μl of the fusion reactions were transferred into a clear-bottom 96-well plate and mixed with 142 μl of nitrocefin developing buffer (100 mM NaPi pH 7.0, 150 μM nitrocefin, 0.2 % Triton X-100). To measure nitrocefin hydrolysis, absorbance at 492 nm was monitored at 15 seconds intervals for 15 minutes at 30°C with a Synergy H1 multimode plate reading spectrophotometer (Biotek, Winooski, VT, USA). Slopes were calculated, and one fusion unit is defined as 1 nmol of hydrolyzed nitrocefin per minute from 12 μg of organelle protein. To block fusion, either antibodies raised against Sec17, Ypt7, Vps33, Vps41, Vam3, Nyv1 or Pep12, or purified recombinant Gdi1 or Gyp1-46 proteins were added to fusion reactions. To drive fusion in the absence of ATP, reactions were supplemented with increasing concentrations of rVam7 in the presence of 10 μg/ml bovine serum albumin. Where indicated, vacuoles were pretreated with 0.2 mM GTPγS for 10 minutes at 27°C prior to addition to fusion reactions. Experimental results shown are calculated from at least 3 biological replicates, each repeated twice (6 technical replicates total).

### Trans-SNARE pairing assay

Trans-SNARE complex formation was assessed as previously described (Schwartz and Merz, 2007). In brief, 45 μg of organelles isolated from BJ3505-CBP-Pep12 *nyv1Δ* (CBY224) or BJ3505-CBP-Vam3 *nyv1Δ* cells and wild type BJ3505 cells were incubated at 27˚C in standard fusion buffer for 60 minutes. Reactions were then placed on ice for 5 minutes and samples, centrifuged (11,000 g, 15 minutes, 4°C), and membrane fractions were resuspended in 560 μl solubilization buffer (20 mM Tris-Cl pH 7.5, 150 mM NaCl, 1 mM MgCl_2_ 0.5% Nonidet P-40 alternative, 10% glycerol) with protease inhibitors (0.46 μg/ml leupeptin, 3.5 μg/ml pepstatin, 2.4 μg/ml pefabloc, 1 mM PMSF). Samples were nutated for 20 minutes at 4°C, centrifuged again (16,000 g, 20 minutes, 4°C) and then supernatants were collected. 10 % of the final extracts were reserved as input samples, and the remaining 90% was brought to 2 mM CaCl_2_. CBP-Pep12 or CBP-Vam3 trans-SNARE protein complexes were then recovered by affinity purification using Calmodulin Sepharose 4B beads: Extracts were added to a slurry of washed beads, nutated overnight at 4°C, collected by brief centrifugation (4,000 g, 2 minutes, 4°C), and washed five times with sample buffer. Beads were then sedimented and bound proteins were eluted by boiling beads (95°C, 10 minutes) in SDS sample buffer containing 5 mM EGTA. Eluates were then used for SDS-PAGE analysis and immunoblotting.

### Western blot analysis

Sodium dodecyl sulfate-polyacrylamide gel electrophoresis (SDS-PAGE) was performed and after separation, proteins were transferred onto a nitrocellulose membrane by wet transfer method (12 V for 8 hours) using a Royal Genie Blotter apparatus (Idea Scientific, Minneapolis, MN, USA). Membranes were blocked with 3% BSA in PBST buffer (137 mM NaCl, 2.7 mM KCl, 10 mM Na_2_HPO_4_, 2 mM KH_2_PO_4_, 0.1% Tween-20) and then washed twice with PBST and incubated with primary antibody diluted to 1:1,000 in PBST for 1 hour at room temperature. Membranes were washed with PBST five times, and then incubated with HRP or FITC labeled goat anti-rabbit or anti-mouse IgG diluted 1:10,000 in PBST for 45 minutes at room temperature. After an additional 5 washes with PBST, the membranes were probed to detect bound secondary antibody using GE Amersham Imager 600 for chemiluminescence or a Typhoon scanner for fluorescence (GE Healthcare, Missassauga, ON, Canada). Experimental results (blots) shown are best representatives of at least 3 biological replicates, each repeated twice (6 technical replicates total).

### Fluorescence microscopy

Live yeast cells were stained with FM4-64 to label vacuole membranes using a pulse-chase method as previously described (Brett et al., 2008). For *in vitro* imaging, organelles isolated from strains expressing Pep12-pHluorin or CPY50-GFP were stained with 3 μM FM4-64 for 10 minutes at 27°C to label vacuole membranes. Micrographs shown were acquired using a Nikon Eclipse TiE inverted microscope equipped with a motorized laser TIRF illumination unit, Photometrics Evolve 512 EM-CCD camera, an ApoTIRF 1.49 NA 100x objective lens, and bright (50 mW) blue and green solid-state lasers operated with Nikon Elements software (housed in the Centre for Microscopy and Cellular Imaging at Concordia University). Micrographs shown are best representatives of at least 3 biological replicates, imaged at least 5 times each (a total of 15 technical replicates) whereby each field examined contained greater than 30 cells or isolated organelles.

### Transmission Electron Microscopy

Isolated organelles were processed for transmission electron microscopy (TEM) as previously described (Mattie et al., 2017). In brief, fusion reactions were incubated at 27°C for 30 minutes, organelles were gently pelleted (5,000 g for 5 minutes) at 4°C and immediately fixed with 2.5% glutaraldehyde in 0.1 M cacodylate buffer (pH 7.4) overnight at 4°C. Membrane pellets then were washed with 0.1 M sodium cacodylate (3 times, 10 minutes) and fixed with 1% osmium tetroxide for two hours at 4°C. Pellets were washed with water (3 times, 5 minutes) followed by gradual dehydration in ethanol (30-100%) and 100% propylene oxide. Pellets were infiltrated with epon:propylene oxide for 1 hour and then embedded in pure epon by polymerization (48 hours at 57°C). Samples were cut into 100 nm thick sections using an ultra-sharp diamond knife and Reichert Ultracut II microtome, loaded onto copper grids, and stained with uranyl acetate (8 minutes) and Reynold's lead (5 minutes). Sections were imaged at 120 kV using an FEI Tecnai 12 electron microscope outfitted with a Gatan Bioscan digital camera (1k × 1k pixels) housed in the Facility for Electron Microscopy Research at McGill University (Montreal, QC, Canada). The micrograph shown is the best representative of 3 biological replicates, imaged at least 5 times (a total of 15 technical replicates) whereby each field contained 4 – 100 organelles depending on magnification used.

### Data analysis and presentation

All quantitative data were processed using Microsoft Excel v.14.0.2 software (Microsoft Cooperation, Redmond, WA, USA), including calculation of means, S.E.M.s and P-values from Student t-tests. n-values shown represent biological replicates. Data were plotted using Kaleida Graph v.4.0 software (Synergy Software, Reading, PA, USA). Micrographs were processed using ImageJ software (National Institutes of Health, Bethesda, MD, USA) and Adobe Photoshop CC (Adobe Systems, San Jose, CA, USA). Images shown were adjusted for brightness and contrast, inverted and sharpened with an unsharp masking filter. All figures were prepared using Adobe Illustrator CC software (Adobe Systems, San Jose, CA, USA).

## AUTHOR CONTRIBUTIONS

C.L.B and M.A.K. conceived the project. S.M. acquired the transmission electron micrograph shown in Figure 1D. D.R.S. acquired fusion data shown in Figures 3E, 4C and 4D. M.A.K. performed all other experiments. M.A.K., D.R.S and C.L.B. prepared data for publication. M.A.K. and C.L.B. wrote the paper.

## ACKNOWLEDGEMENTS

We thank J.E. Gerst, A.J. Merz and W.T. Wickner for custom antibodies, yeast strains and plasmids, as well as C. van Oostende-Triplet and C. Law for technical assistance with microscopy. The Canadian Foundation for Innovation and Natural Sciences and Engineering Research Council of Canada (NSERC) provided generous support for the Centre for Microscopy and Cell Imaging at Concordia University. D.R.S. is a postdoctoral scholar funded by the Olle Engkvist Byggmästare Foundation. This research was funded by Concordia University and by grants RGPIN/403537-2011 and RGPIN/2017-06652 to C.L.B from the NSERC Discovery program.

## LIST OF ABBREVIATIONS

ALP: Alkaline Phosphatase;
BLOC: Biogenesis of Lysosome-like Organelles Complex-1;
CPS: CarboxyPeptidase-S;
CPY: CarboxyPeptidase-Y;
ESCRT: Endosomal Sorting Complexes Required for Transport;
GAP: Gtpase Activating Protein;
GEF: Guanine nucleotide Exchange Factor;
HOPS: HOmotypic fusion and vacuole Protein Sorting;
ILF: IntraLumenal Fragment;
ILV: IntraLumenal Vesicle;
MTC: Multisubunit Tethering Complex;
MVB: MultiVesicular Body;
NSF: N-ethylmalemide-Sensitive Factor;
SNAP: Soluble Nsf Attachment Protein;
SNARE: SNAp REceptor;
TEM: Transmission Electron Microscopy;
VPS: Vacuole Protein Sorting.

## LIST OF SUPPLEMENTAL MATERIAL

Figure S1, related to Figure 1. Organelle isolation by sucrose gradient does not permit fusion Figure S2, related to Figure 3. Fusion probes are properly localized in *nyv1*∆ cells

---

**Figure S1.**
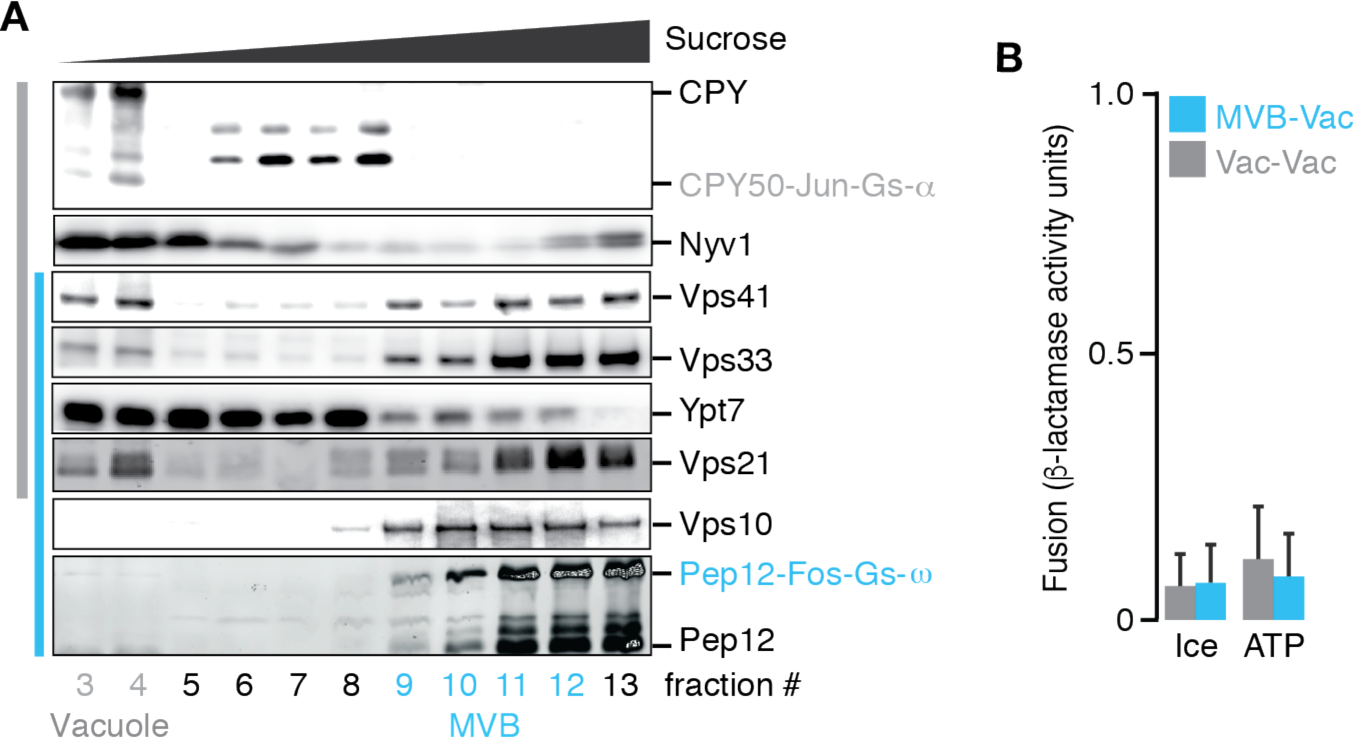
Organelle isolation by sucrose gradient does not permit fusion. (**A**) Western blots to confirm the presence of MVB (Vps10) and vacuole (CPY) markers, fusion probes and fusogenic proteins in organelles isolated by sucrose gradient. (**B**) Content mixing values obtained after isolated organelles were incubated for 90 minutes at 27˚C or placed on ice (negative control) in the presence of ATP to trigger fusion. Reactions contained fractions 9-12 (MVBs) mixed with fractions 3-4 (vacuoles) isolated from separate strains expressing complimentary fusion probes targeted to MVBs or vacuoles (MVB-Vac), or probes only targeted to vacuoles (Vac-Vac). Means ± S.E.M shown (n ≥ 3).

**Figure S2.**
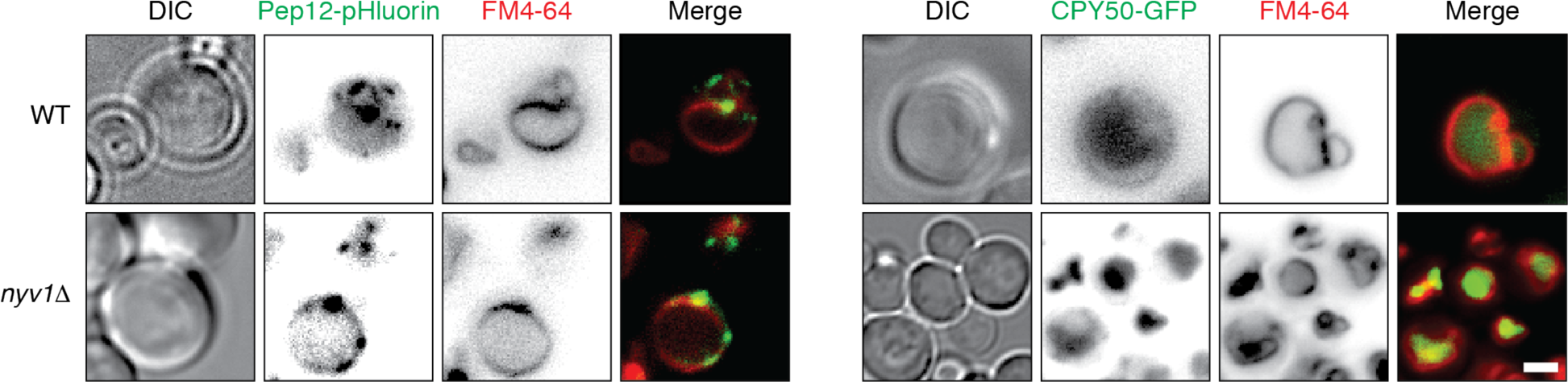
Fusion probes are properly localized in *nyv1*∆ cells. Fluorescence micrographs of live wild type (WT) or *nyv1*∆ cells expressing Pep12-pHluorin (left) or CPY50-GFP (right). Vacuole membranes are stained with FM4-64. Scale bar, 2 μm.

